# In silico identification of GPI-anchored proteins in Paracoccidioides

**DOI:** 10.1101/347351

**Authors:** L.R. Basso, R.A. Gonçales, E.J.R Vasconcelos, T.F. Reis, P. C. Ruy, J.C. Ruiz, P.S.R. Coelho

**Author notes:** These authors contributed equally to this work (L.R. Basso, R.A. Gonçales). Mailing address: Department of Cellular and Molecular Biology and Pathogenic Bioagents, Faculty of Medicine of Ribeirão Preto – University of São Paulo – USP, Avenida dos Bandeirantes, 3900, Ribeirão Preto, SP 14049-900.

## Abstract

Glycosylphosphatidylinositol-anchored proteins (GPI-proteins) are widely found in eukaryotic organisms. In fungi, GPI-proteins are thought to be involved in diverse cellular mechanisms such as cell wall biosynthesis and cell wall remodeling, adhesion, antigenicity, and virulence. The conserved structural domains of GPI-protein allow the utilization of *in silico* prediction approach to identify this class of proteins using a genome-wide analysis. We used different previously characterized algorithms to search for genes that encode predicted GPI-proteins in the genome of *P. brasiliensis and P. lutzii*, thermal dimorphic fungi that causes paracoccidioidomycosis (PCM). By using these methods, 98 GPI-proteins were found in *P. brasiliensis* with orthologs in *P. lutzii*. A series of 28 GPI-proteins were classified in functional categories (such as glycoside hydrolases, chitin-processing proteins, and proteins involved in the biogenesis of the cell wall). Furthermore, 70 GPI-proteins exhibited homology with hypothetical conserved proteins of unknown function. These data will be an important resource for the future analysis of GPI-proteins in *Paracoccidioides spp*.

## 1. Introduction

In pathogenic fungi, cell wall components mediate the contact, adhesion, and invasion of the host tissue (Levitz, 2010) which is characterized by a distinctive composition depending on the morphological phase. The covalently linked glycosylphosphatidylinositol (GPI)-proteins represents the main class of proteins in the cell wall. Members of this group are pivotal players in cell wall biogenesis and remodeling and in essential pathogenic processes such as adhesion to and degradation of host tissues (Hube and Naglik, 2001; Martchenko et al., 2004; Richard et al., 2002; Richard and Plaine, 2007; Sundstrom, 2002). All GPI-proteins show common characteristics such as a C-terminal consensus sequence for GPI modification and an N-terminal signal peptide for translocation across the membrane of the endoplasmic reticulum. In addition, many GPI-proteins are heavily glycosylated. These well-defined characteristics allow the use of *in silico* approaches for screening and identification of GPI-proteins in the genome (De Groot et al., 2003; Eisenhaber et al., 2004).

The *Paracoccidioides* genus includes species of thermodimorphic pathogenic fungi that cause paracoccidioidomycosis (PCM) which is the most prevalent systemic human mycosis occurring in Latin America. Multilocus and genomic sequencing studies indicate the existence of two distinct species within the *Paracoccidioides* genus, *P. brasiliensis* and *P. lutzii* (4). *P. brasiliensis* is comprised of distinct lineages (S1, PS2, PS3, and PS4) that cause most of PCM cases throughout South America (4–6). *Paracoccidioides lutzii* isolates are found predominantly in north, central and southwest of Brazil and Ecuador (Teixeira et al., 2009; Theodoro et al., 2012).

The complete genome sequence of three strains (Pb18, Pb03 and Pb01-like) from the *Paracoccidioides* genus (Desjardins et al., 2011a; Munoz et al., 2014) have been recently available. The genomic information has allowed the identification, in *P. brasiliensis and P. lutzii*, of a group of 62 predicted GPI-proteins based on an *in silico* analysis using one program of prediction (Desjardins et al., 2011a, b). Based in the current genome sequence *of P. brasiliensis* and *P. lutzii* we used five algorithms to look for additional GPI-proteins that may not have been detected by the previous genome analysis. The identification of new *P. brasiliensis* GPI-proteins may reveal targets to be used in the future for the development of new diagnostic tools, drugs, and immunologic tests.

## 2. Material and Methods

### In silico identification of predicted GPI-proteins and transcription analysis

The pipeline for the identification of GPI-protein coding genes consisted of the following steps: (1) A proteome file containing 8741 ORF sequences of *P. brasiliensis* strain Pb18 and 9.132 ORF sequences of *P. lutzii* (Desjardins et al., 2011a, b) were downloaded from the Broad Institute website (http://www.broad.mit.edu) last accessed on March 2013. These proteomes were searched for proteins carrying a GPI anchor signal peptide (omega site) using five different methods: one developed by de Groot and collaborators (De Groot et al., 2003), Big-PI (http:// Eisenhaber et al., 2004), GPI-SOM (http:// Fankhauser and Maser, 2005), PredGPI (http://Pierleoni et al., 2008a, b) and FragAnchor (http:// Poisson et al., 2007). (2) predicted-GPI proteins detected by at least two methods were selected for further analysis. Protein sequences from *P. brasiliensis* and *P. lutzii* that presented mismatches after alignment by ClustalW were reannotated (see the following section) (3) Proteins were searched for the presence of an N-terminal ER-import signal by SignalP V3.0 software available at (http://www.cbs.dtu.dk/services/SignalP-3.0). The standardized threshold value of 0.5 was applied for signal peptides in the two algorithms [SignalP-NN (Smean) and SignalP-HMM (Sprob)]. (4) The presence of internal membrane domains and protein localization was investigated using TMHMM (http://www.cbs.dtu.dk/services/TMHMM/) and PSORTII (http://psort.nibb.ac.jp).

Predicted GPI proteins were used as queries in BLASTp searches (Altschul et al., 1990) to identify sequence similarities in 17 other fungal genomes (Cuomo and Birren, 2010). After the alignment with *ClustalW* (http://www.genome.jp/tools/clustalw), we considered orthologues when their protein sequence presents > 30 % of amino acid identity in a region encompassing > 60 % of the whole sequence. BLASTP searches were performed at NCBI (http://www.ncbi.nlm.nih.gov/blast).

For transcription analysis, ESTs were manually searched for hits in the Broad Institute database.

### Annotation and manual curation of GPI-protein coding genes

Genomic DNA sequences encompassing ORFs coding for GPI-anchoring signals – from 1000 bp upstream the start codon to 1000 bp downstream the stop codon - were analysed by five programs to determine the exon-intron map: *Augustus* (http://bioinf.uni-greifswald.de/augustus/), *Fgenesh* (http://linux1.softberry.com), *GeneMark* (http://opa.biology.gatech.edu/GeneMark/, *Geneid* (http://genome.crg.es/geneid.html) and *Genscan* (http://genes.mit.edu/GENSCAN.html),. The consensus gene sequence from all prediction strategies was aligned by *ClustalW* with the orthologues of *P. brasiliensis* (Pb18, Pb03) and *P. lutzii* (Pb01) whose sequences are from the NCBI fungal data bank. The annotated sequences were re-analyzed to confirm the presence of GPI-anchoring signal and signal peptide, and absence of transmembrane domains as described in the previous section.

## 3. Results and Discussion

### *In silico* identification of GPI proteins in *P. brasiliensis*

Five different *in silico* methods were used to comprehensively identify putative GPI proteins in the *P. brasiliensis and P. lutizi* genome (De Groot et al., 2003; Eisenhaber et al., 2004; Fankhauser and Maser, 2005; Pierleoni et al., 2008a; Poisson et al., 2007). The resulting genomic sequences were reannotaded when there was a discrepancy in the exon-map structure after alignment with conserved genomic sequences from close relatives (e.g *P. brasiliensis* X *P. lutzii*; see materials and methods and supplementary data). We identified 28 GPI-proteins that can be classified into 5 functional categories, 70 conserved proteins with unknown functions, in a total of 98 GPI-proteins (Table 1). The ratio of predicted GPI-proteins (1.12%) to the proteome is similar to that observed in filamentous fungi and yeasts (de Groot et al., 2009; de Groot et al., 2003; Desjardins et al., 2011; Eisenhaber et al., 2004). We did not find differences in the number of GPI-proteins homologs between *P. brasiliensis* and P. *lutzii*.

**Table 1.** Predicted GPI proteins of *Paracoccidioides*.

| *P. brasiliensis*<br>ORF ID | *P. lutzii*<br>ORF ID | Size (aa -<br>Pb/PI) | Identity<br>(%) | Big-PI<br>(Pb/PI) | FragAnchor<br>(Pb/PI) | PredGPI<br>(Pb/PI) | DeGroot<br>(Pb/PI) | GPI-SOM<br>(Pb/PI) | SignalP<br>(Pb/PI) | EST | function/family |
| --- | --- | --- | --- | --- | --- | --- | --- | --- | --- | --- | --- |
| **Carbohydrate modifying enzymes and resident proteins** |  |  |  |  |  |  |  |  |  |  |  |
| 04499 | 07670 | 370/389 | 95.7 | + | H | H | + | + | -/+ | 45/47 | ECM33 |
| 02460 | 04708 | 225/225 | 79.2 | +/+ | H/H | H/H | +/+ | +/+ | + | 9/7 | PbPga1 |
| 04918 | 05068 | 529/529 | 97.1 | + | H | H | + | + | + | 29 | GH2 |
| 05076 | 05245 | 471/471 | 98.1 | + | H | H | + | + | + | 6 | GH72 (Gel) |
| 04328 | 06867 | 224/224 | 94.6 | + | L/H | W/P | + | + | - | - | GH16 (Crh) |
| 04416 | 06779 | 436/435 | 77.7 | +/- | L/- | H/- | +/- | +/- | + | 5 | GH16 (Crh) |
| 06728 | 05380 | 754/743 | 96.9 | +/- | L/- | H/- | + | + | + | - | GH20 |
| 03691 | 01139 | 432/431 | 92.1 | +/+ | - | H/P | +/+ | +/+ | + | - | GH16 (Crh) |
| 04432 | 06764 | 532/531 | 92.8 | +/- | H/H | P/H | +/+ | +/+ | + | 3 | GH13 (Alpha-amilase) |
| 04693 | 04534 | 459/459 | 97.6 | -/+ | H | -/W | + | + | + | 8 | GH76 (DCW1) |
| 05063 | 05192 | 443/506 | 92.5 | - | P/P | - | - | + | - | - | GT1 |
| 01494 | 06960 | 470/471 | 88.7 | -/- | - | H/H | +/+ | +/+ | + | - | GH76 (DCW1) |
| 03286 | 03782 | 445/443 | 79.4 | -/+ | -/P | H | - | + | + | 3 | GH72 (Gel) |
| 07673 | 05717 | 323/323 | 95.4 | - | - | H/W | - | - | + | - | GH76 (DCW1) |
| 00483* | 01798 | 862/852 | 95.6 | - | - | - | -/- | -/+ | +/- | - | GH3 |
| 00795 | 02100 | 446/446 | 97.3 | - | - | - | + | + | + | 3 | GH72 (Gel) |
| 03601 | 01224 | 265/363 | 25.7 | - | - | - | +/- | - | - | - | GH16 (Crh1) |
| 04617 | 04461* | 234/237 | 93.6 | - | - | - | + | -/+ | +/- | 1 | CE5 (cutinase) |
| 06344 | 03482 | 796/785 | 96.5 | - | - | - | + | - | + | 5 | GH92 |
| 06465 | 03359 | 230/230 | 90.0 | - | - | - | + | - | + | - | GH23 (lysozyme) |
| **Aspartic proteases** |  |  |  |  |  |  |  |  |  |  |  |
| 06207 | 03619 | 513/469 | 70.4 | -/+ | -/P | -/H | -/+ | -/+ | + | 2 | Pepsin-like proteinase |
| 08282 | 07583 | 481/481 | 95.6 | +/- | P/HP | P/HP | +/+ | +/+ | - | 2 | Pepsin-like protease |
| **Phospholipases** |  |  |  |  |  |  |  |  |  |  |  |
| 01064 | 06610* | 1067/1004 | 96.3 | - | - | - | + | + | - | 7/9 | Phospholipase D<br>Active site |
| 03299 | 03796 | 1424/1398 | 93.3 | - | - | - | + | + | + | 4/4 | phospholipase C X<br>domain |
| **Hydrophobins** |  |  |  |  |  |  |  |  |  |  |  |
| 08524 | - | 419 |  | + | L | H | + | + | + | - | Hydrophobin |
| 07472 | 04235* | 133/133 | 100 | - | - | P/P | + | - | +/+ | 2829 | Hydrophobin 2 |

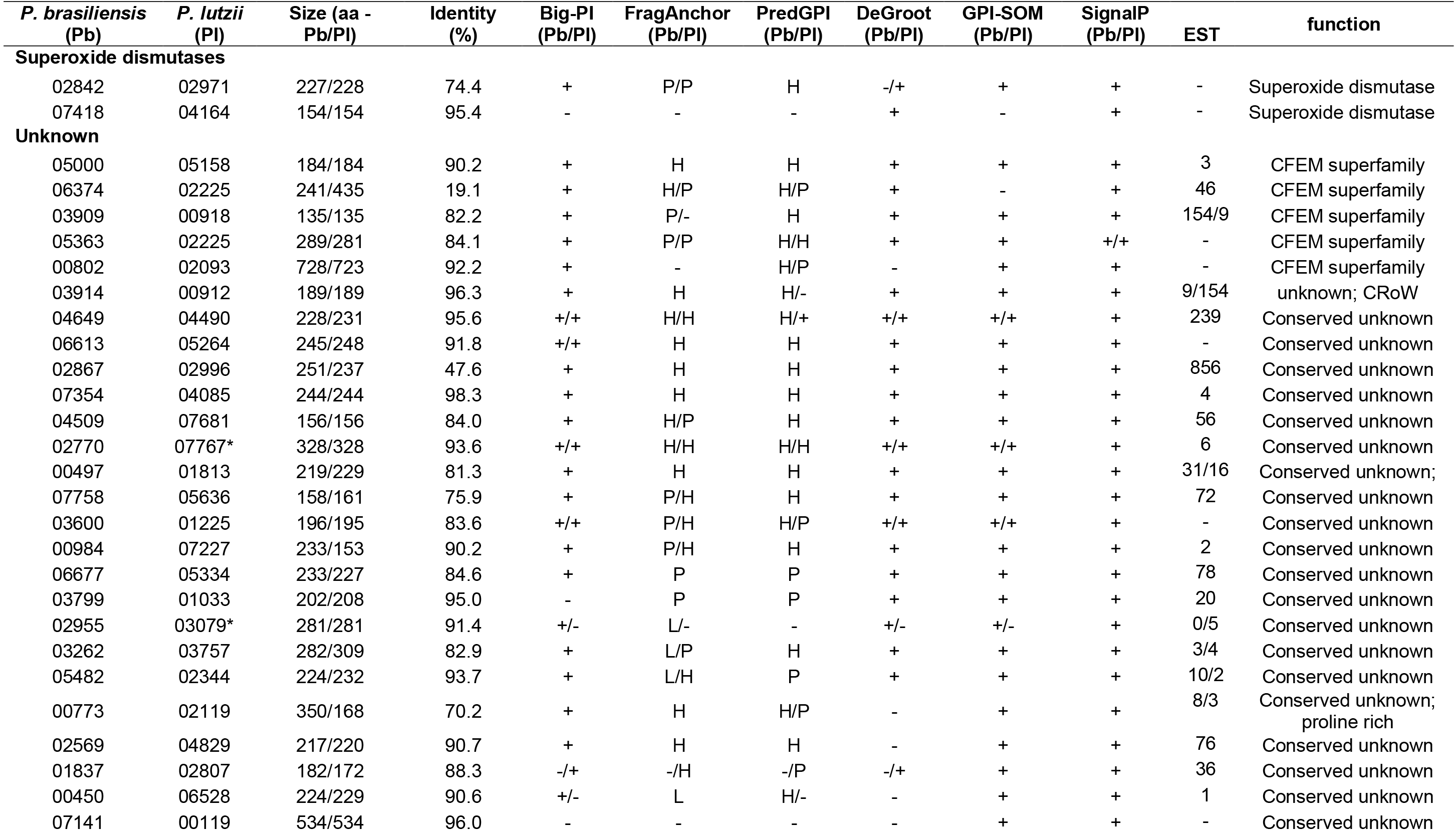

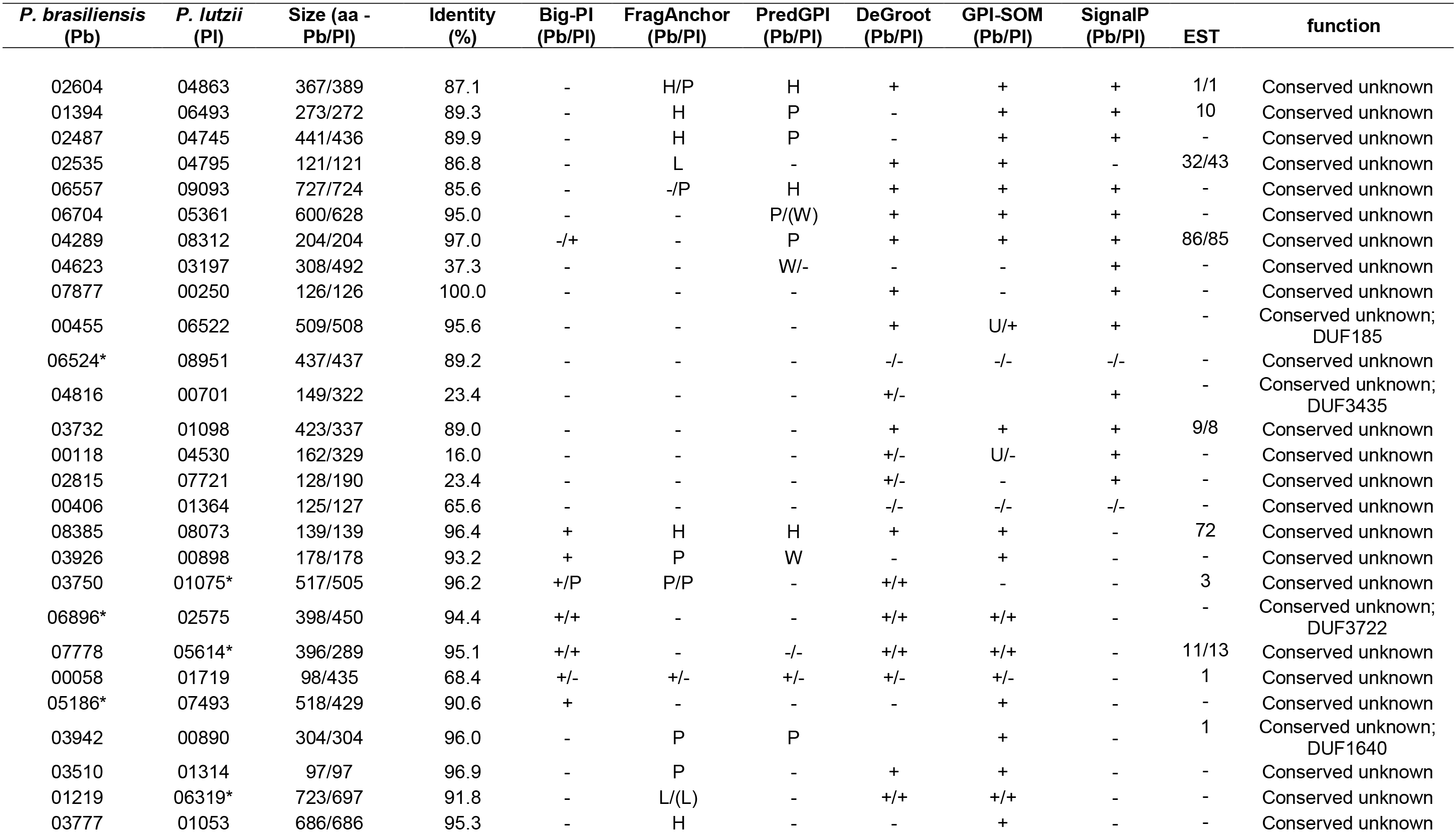

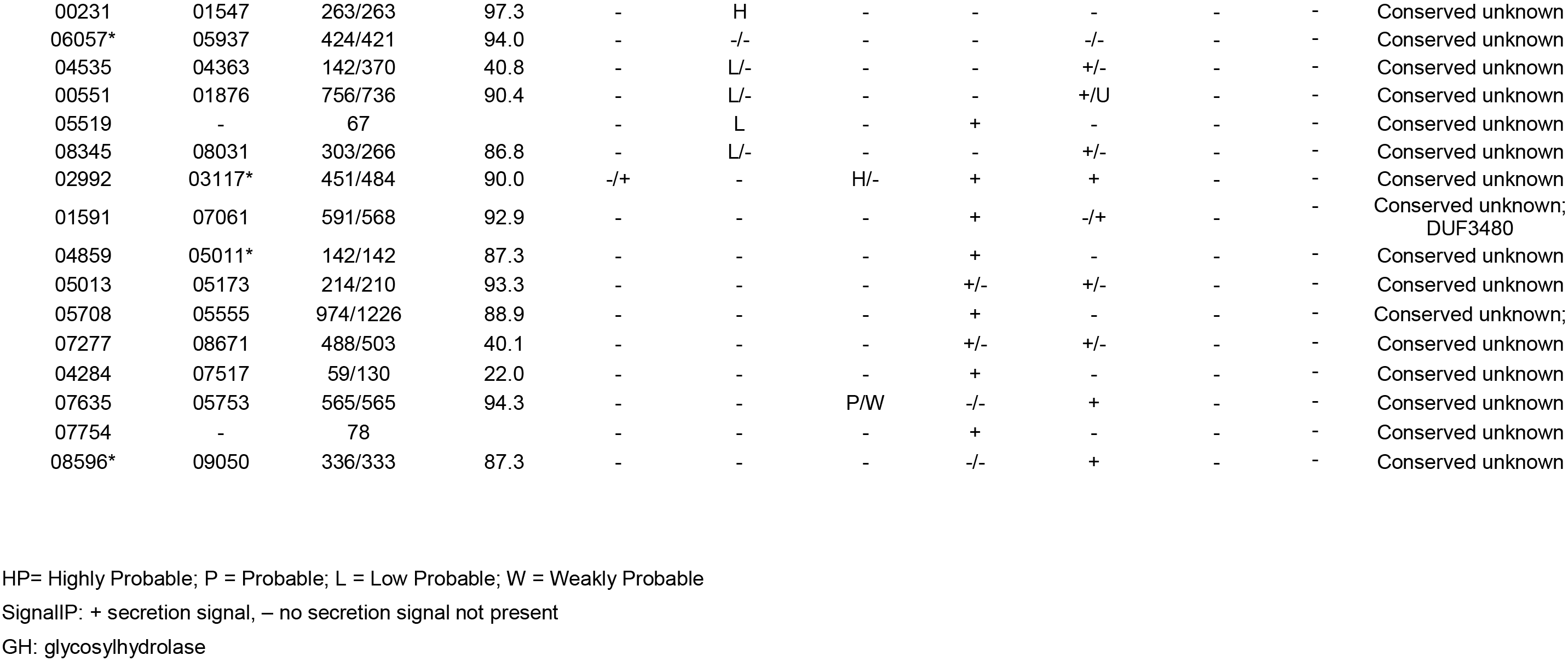

### Synthesis and processing of glycan, chitin and other enzymatic processes of the cell wall

Sixteen of the predicted GPI-proteins have homology with glycosyl hydrolases (GH), (Cantarel et al., 2009; Lairson et al., 2008). Three proteins belong to the GH72 family or Gel/Gas/Phr of 1,3–β- glucanosyltransferases (Castro Nda et al., 2009; Mouyna et al., 2000; Popolo et al., 2008). Four GPI-proteins belong to the Crh (GH16) family which consists mainly of 1,3–β-glucanases and 1,3-1,4–β- glucanosyltransferases (Cabib et al., 2008). Three members of the Dfg5 (GH76) family were also identified, and they are endomananases with a putative role in the incorporation of GPI-proteins to the cellular wall (Maddi et al., 2012). An alfa-amylase (GH13) is present and has a putative role of in the processing of 1,3-α-glucana/1,4-α-glucana (Camacho et al., 2012). The PbPga1 is a cell wall protein with immunogenic properties and is recognized specifically by serum of patients with paracoccidioidomycosis (Valim et al., 2012).

The presence of a lysozyme (GH23) is interesting. The biological function of this enzyme in fungi has not yet been determined. It can be speculated that the molecule can serve as a tool in the breakage of the bacterial peptidoglycan for nutritional purposes (Korczynska et al., 2010). An N-acetyl-β-D-glicosaminidase (GH20) was identified (Santos et al., 2004) and is a member of the GH92 family. Finally, it was identified a member of the GH3 family, a glucosyl-transferase (GT1) and a cutinase (CE5). Cutinases are extracellular enzymes secreted by microorganisms to break the cellular wall of plants (Lin and Kolattukudy, 1980). The protein is possibly a false positive because these enzymes are secreted (de Vries, 2003).

### Cell wall biogenesis

One of the proteins identified (Ecm33p) is thought to play an important role in cell wall biogenesis. This protein shares homology with the Ecm33p family, also a ubiquitous fungal GPI protein family. Deletion of *ECM33* in *S. cerevisiae* results in swollen cells that secrete increased levels of 1,6-β-glucosylated CWPs into the culture medium and are hypersensitive to drugs that interfere with cell wall biosynthesis (De Groot *et al.*, 2001).

### Proteins with varied functions

Four proteins share homology with fungal proteins not directly associated with the cell wall. We identified two superoxide dismutases (SOD). SOD converts superoxide radicals into hydrogen peroxide and molecular oxygen and is thought to be important to the pathogenesis of fungal infection. Two proteins share homology with hydrophobins, which in filamentous fungi play a role in the formation of aerial hyphae and in the attachment of hyphae to hydrophobic surfaces. The protein PbPga1 was first identified by homology with a predicted GPI-protein of *Histoplasma capsulatum*. PbPga1 has been characterized as a cell surface protein recognized by sera from patients with paracoccidioidomycosis (Valim, 2012)

### Conservation of *Paracoccidioides brasiliensis* GPI-proteins

We analyzed the conservation of predicted GPI proteins of known function in 17 different fungal genomes. These proteins represent the most conserved proteins (Fig 1) and are involved in different biosynthetic and processing pathways (glycosylases, proteases and lipases). On the other hand, the less conserved proteins (Fig 2) are those with unannotated function. These proteins may not have catalytic roles in the cell walls. Many of the identified proteins with unknown function are expressed as evidenced by EST hits in databases (Table 1)

**Fig 1.**
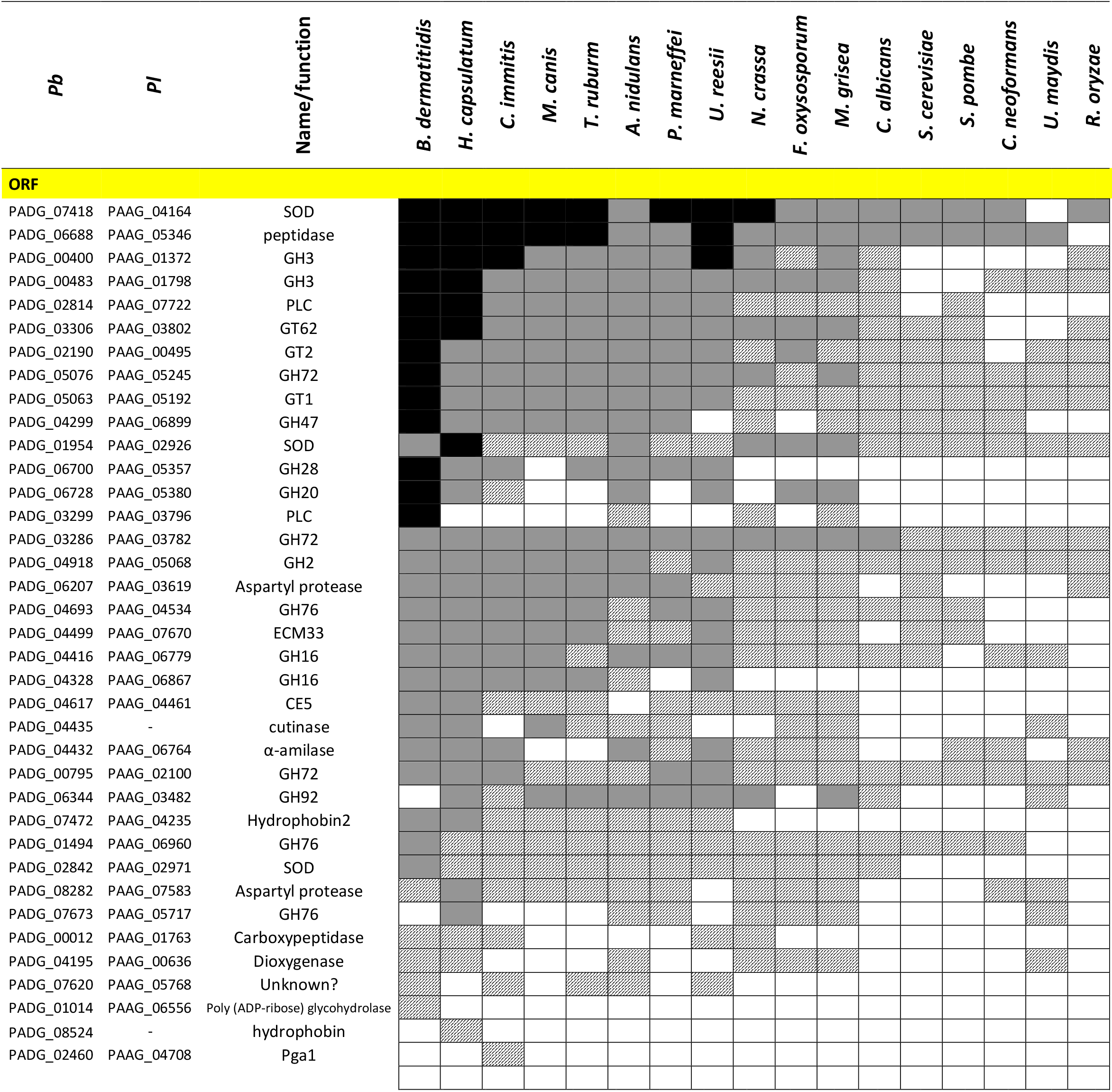
Presence of homologs for GPI proteins with known function. GT62 = α-1-6 mannosyltransferase GT2 = dolichyl-phosphate beta-glucosytransferase

**Fig 1.**
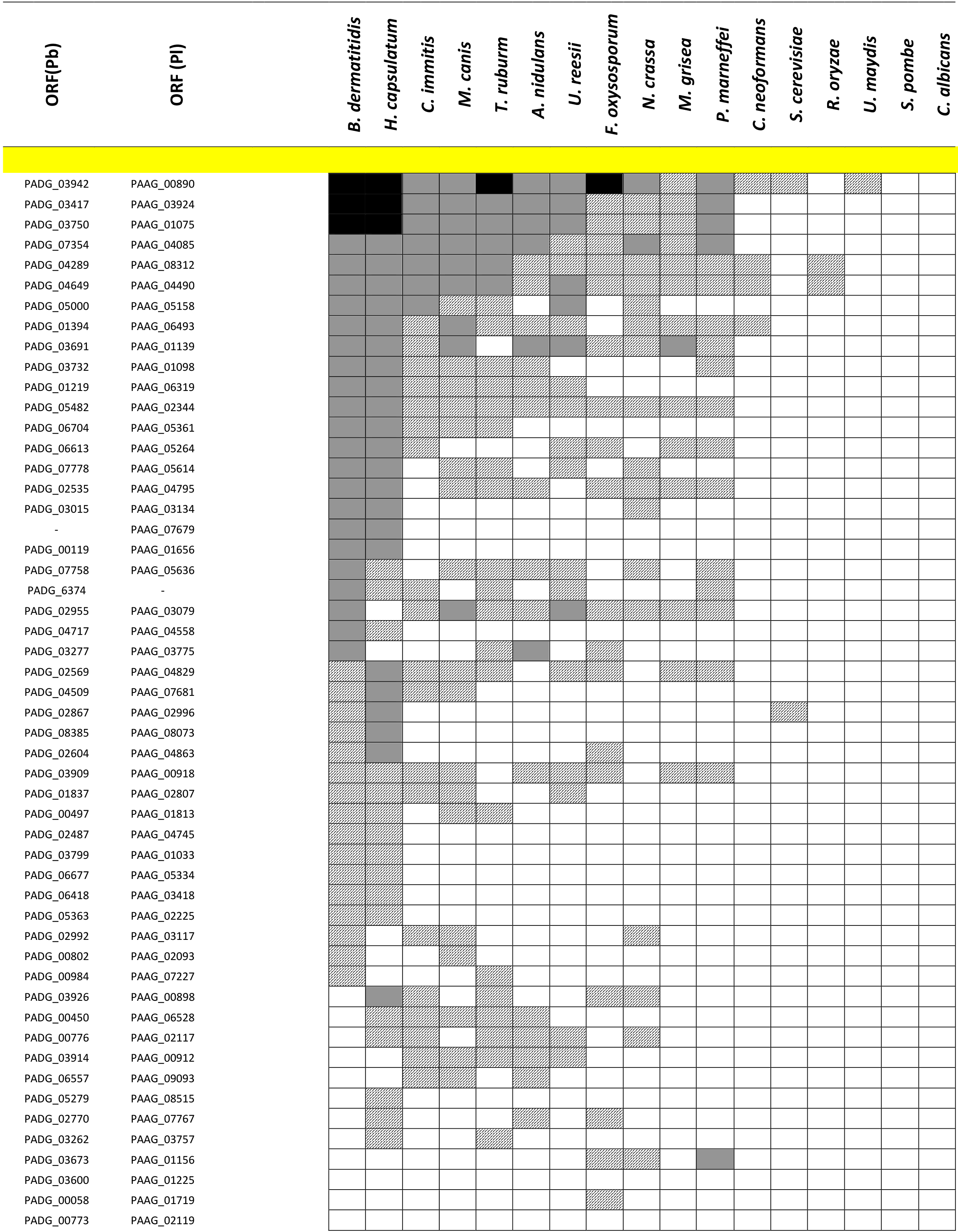
Presence of homologs for GPI proteins with unknown function.

